# First record of double-edged sword effect of caterpillar-induced plant volatiles in nature

**DOI:** 10.1101/093963

**Authors:** Ashraf M. El-Sayed, David M. Suckling

## Abstract

Plants release volatiles in response to caterpillar feeding that attract natural enemies of the herbivores, a tri-trophic interaction which has been considered an indirect plant defence against herbivores. The caterpillar-induced plant volatiles have been reported to repel or attract conspecific adult herbivores. Apple seedlings infested with *Pandemis pyrusana* larvae uniquely release five compounds (benzyl alcohol, benzyl nitrile, phenylacetaldehyde, indole, and (*E*)-nerolidol). These compounds and other known caterpillar-induced plant volatiles were tested to investigate the response of both herbivores and natural enemies. In field tests, binary blends of benzyl nitrile and acetic acid or 2-phenylethanol and acetic acid attracted a large number of conspecific male and female adult moths. On the other hand, a ternary blend of benzyl nitrile, 2-phenylethanol and acetic acid attracted the largest numbers of the general predator, the common green lacewing, *Chrysoperla carnea.* This study provides the first record of caterpillar-induced plant volatile attraction to conspecific adult herbivores as well as predators under natural conditions.

## Introduction

Insect herbivory typically induces a change in the profile of volatile organic compounds released by the affected plants (Dicke and Baldwin 2010; Hare JD. 2011). Natural enemies of insect herbivores such as predators and parasitoids are attracted to these altered plant odours and as a consequence are better able to locate their herbivore hosts (Turlings et al. 1990; Alborn et al. 1997; Yoshinaga et al. 2010). This tri-trophic relationship is considered an indirect plant defence strategy that recruits natural enemies that incapacitate the herbivores, resulting in higher plant fitness. In contrast, feeding by caterpillars on plants can result in contrasting behaviour by conspecific adult herbivores, in some plant species, volatiles induced by caterpillar feeding repel conspecific adult herbivores e.g. (De Moraes et al. 2001; Signoretti et al. 2012); while odours of other plant species infested with caterpillars can attract conspecific adults e.g. (Rojas 1999; Sun et al. 2014). This suggests two different paradigms for the role of herbivore-induced plant volatile compounds (HIPVC). In the first paradigm, HIPVC attract natural enemies and repel herbivores (single-edged sword effect), while in the second paradigm, HIPVC attract both natural enemies and herbivores (double-edged sword effect). The advantages in the first paradigm are theoretically sound because the ability of adult herbivores to detect and avoid oviposition on damaged plants would have several advantages including herbivore offspring avoiding competition for food resources, reducing the probability of encountering natural enemies, and avoiding increased host plant resistance with lower nutritional content (De Moraes et al. 2001). The discrepancies reported in several plant-insect systems in relation to the response of herbivores to HIPVC could simply be due to experimental problems and biased toward demonstrating the beneficial effects of HIPVC as an indirect plant defence strategy. In addition, most of these studies were designed to investigate individual response in multi-tropic systems under laboratory conditions, rather than focusing on the collective responses of herbivores and natural enemies under natural conditions.

Among the major tortricid pests in North American orchards are a suite of leafrollers that cause significant losses in fruit production. This includes the oblique-banded leafroller (OBLR), *Choristoneura rosaceana* (Harris), the eye-spotted bud moth (ESBM), *Spilonota ocellana* (Denis & Schiffermüller), and the Pandemis leafroller (PLR), *Pandemis pyrusana* (Kearfott). The PLR has a wide host range and occur in apple and pear orchards. These species have been considered important pests of apples in the western US and Canada (Deland et al. 1994). Understanding the tri-trophic relationship between host plants and both herbivores and natural enemies would enable a more effective semiochemical-based system to control these pests.

Apple seedlings uniquely released several compounds including acetic acid, acetic anhydride, benzyl alcohol, benzyl nitrile, indole, 2-phenylethanol, and (E)-nerolidol only when infested by larvae of light brown apple moth (LBAM), *Epiphyas postvittana* (Walker) (Suckling et al. 2012; El-Sayed et al. 2016). Recently, El-Sayed et al. 2016 found that a blend of the two HIPVC, benzyl nitrile and acetic acid attracted a significant number of conspecific male and female adult LBAM in New Zealand. Further investigation with other leafrollers (Tortricidae) in North America including the ESBM and OBLR revealed similar systems. Male and female adults of OBLR were most attracted to a blend of 2-phenylethanol and acetic acid. Our counter-intuitive results described in (El-Sayed et al. 2016) are the first identification of caterpillar-induced plant volatiles that attract (or repel) insect herbivores. The demonstration of biological activity of HIPVC in the attraction of conspecific adults in three tortricid species in two biogeographic regions different from the origin of the apple *(Malus)* suggests a widespread phenomenon. However, in the previous study, we did not report the response of natural enemies to HIPVC.

The present work was undertaken to investigate the response of apple trees to infestation with Pandemis leafroller, *P. pyrusana* larvae and to investigate the collective response of both adult herbivores and natural enemies to HIPVC under natural conditions, which is generally lacking from the literature. This study was expected to be a test case for either the double-edged sword effect hypothesis (attraction of HIPVC to both natural enemies and herbivore), or single-edged sword effect hypothesis (attraction of HIPVC to natural enemies and repellence to herbivore).

## Materials and methods

### Plants and Insects

Pandemis leafrollers. *P. pyrusana* (PLR) were obtained from a laboratory colony at Washington State University, Wenatchee, WA that was established in 1985 from larvae collected from Yakima, WA. This colony has been reared continuously since their collection on a pinto bean diet following the method of (Shorey and Hale 1965) under constant conditions of temperature (23 ± 2 °C), relative humidity (RH, 70%), photoperiod (16:8, L:D), and without exposure to insecticides. Neonate 4^th^ instar PLR were transferred with a brush to new shoots on 2-year-old, potted ‘Fuji’ apple trees at the USDA Laboratory in Wapato, WA. Three to five larvae were transferred to each actively-growing shoot on several trees.

### Chemicals

Chemical purity of the standards used to identify the compounds in infested apple seedling headspace and used in the field experiments were as follows: Glacial acetic acid (99%), benzyl alcohol (99%), (E)-nerolidol (85%), benzyl nitrile (99%), phenylacetaldehyde (99%), 2-phenylethanol (99%), and indole (99%). Glacial acetic acid was stored under ambient temperature while all other compounds were stored at −20°C until used. All compounds were purchased from Sigma Aldrich (MO, USA).

### Air Entrainment of Volatiles Emitted by Apple seedlings infested with PLR Larvae

Volatile collections from infested apple trees (cv. Red Jonaprince) with PLR larvae and uninfested apple trees were conducted in Yakima, USA using a dynamic headspace collection method, where air containing the odor was absorbed by a sorbent filter that was then extracted by solvent. Intact tree branches with either apple leaves infested with leafroller larvae or uninfested leaves were enclosed in a polyester oven bag (Glad NZ®, 35 cm × 50 cm). A charcoal-filtered air stream was pulled over the enclosed leaves at 0.5 L/min, and the headspace volatiles were collected for 24 h on an adsorbent filter containing 50 mg of Tenax-GR 35/60 (Alltech Associates Inc.) in a 60 mm long × 6 mm diameter glass tube. For collection of control samples, a charcoal-filtered air stream was pulled through an empty oven bag in the same greenhouse. Samples were sealed and shipped in dry ice to Plant and Food Research (PFR) facility for GC/MS analysis. At the PFR lab, the Tenax filters were extracted with 0.5 ml of n-hexane (AnalaR BDH, Laboratory Supplies, Poole, UK). A sub-sample of 100 μl was reduced to 10 μ1 at ambient temperature under a stream of argon and 1 μl of the concentrated extract was injected in the GC/MS. Six volatile collection samples from infested and uninfested leaves and six control samples were sampled.

### Analysis of Air-Entrainment Samples by Gas Chromatography/Mass Spectrometry (GC/MS)

The concentrated extracts of the air-entrainment samples were analyzed using GC/MS (Varian 3800 GC coupled to a Varian 2200 MS). Helium was used as the carrier gas (1 mL min^−1^), and injections were splitless for 0.6 min. Transfer line and ion trap temperatures were 250 and 180 °C, respectively. The GC injector temperature was set at 220 °C, and the oven ramp was 40 °C for 2 min, 4 °C min^−1^ to 240 °C, hold for 10 min, and then 15 °C min^−1^ to 260 °C, using a VF-5 MS capillary column (30 m × 0.25 mm inner diameter × 0.25 μm film; and a polar 30 m × 0.25 mm i.d. × 0.5 μm, VF23-MS capillary column; Varian, Inc., Walnut Creek, CA). A 1 μL aliquot was injected after first concentrating 100 μL of each sample to ca. 10 μL with a gentle stream of argon. The spectra were recorded at an ionization voltage of 70 eV over a mass range mass-to-charge (*m/z*) of 20 to 499. Kovats retention indexes (KI) were calculated for the compounds (El-Sayed 2016) (Table 1). Structural assignments of the compounds were made by comparing their mass spectra with the MS library (NIST 2002), as well as by comparison to Kovats retention indices published in the literature (El-Sayed 2016). Identification of volatiles was confirmed by comparison to authentic samples.

**Table 1.**
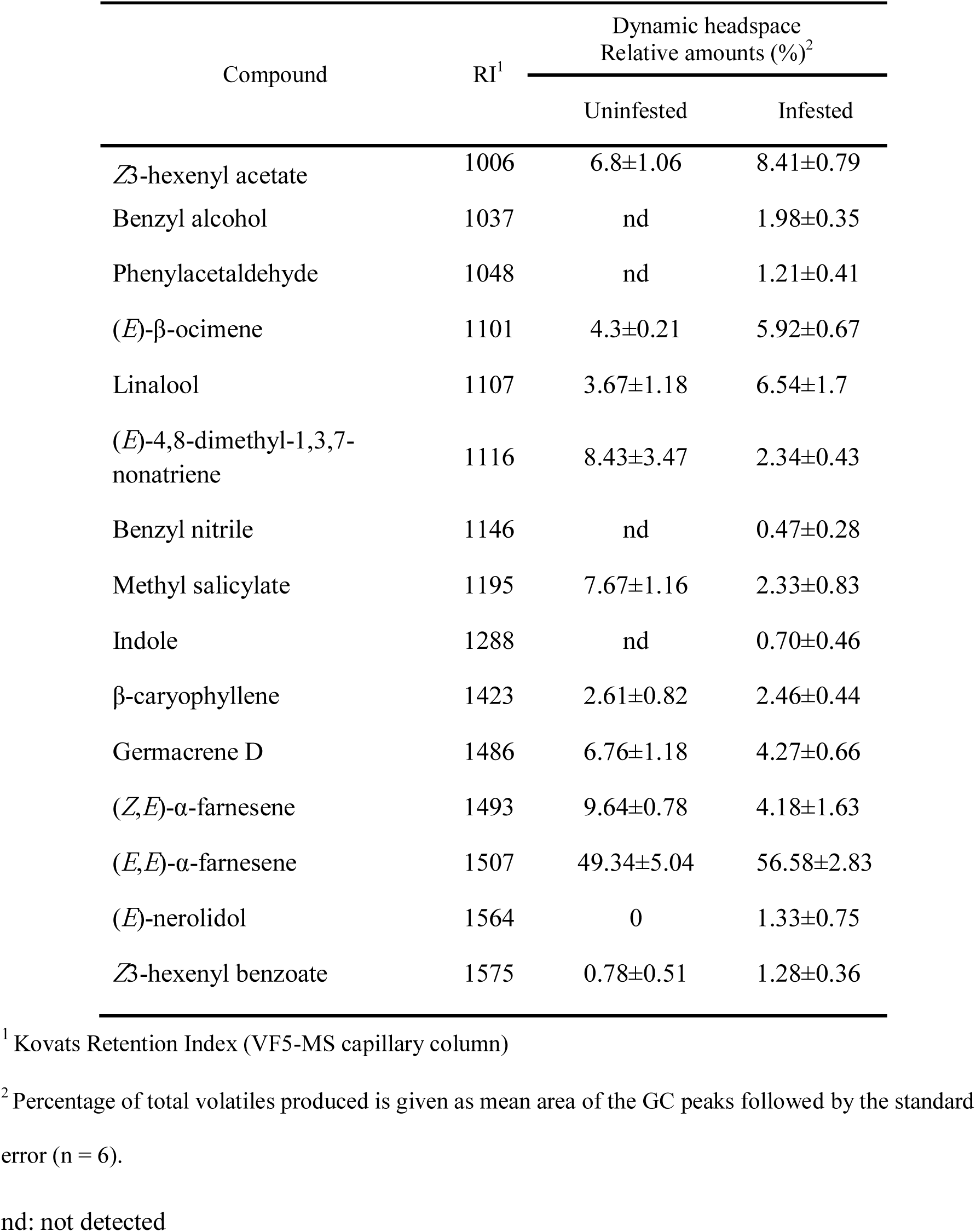
Relative amounts (% ±SEM) of the compounds identified in the headspace of uninfested apple seedlings and apple seedlings infested with *P. pyrusana* larvae.

### Field Experiments

The two field experiments were conducted in a mixed varieties apple orchard in Washington, USA. Large white delta traps (Pherocon VI, Trécé Inc, Adair, OK) were used in the two trials. For each treatment, 100 μ1 of neat chemical were pipetted into a 5 × 5 cm polyethylene sachet with a thickness of 100 μm containing a rectangular piece of wool felt (4 × 2 cm2). Acetic acid dispenser were made by pipetting 3 ml of glacial acetic acid in a 5 ml polyethylene vial with 3 mm bore size in the vial lid (Thermo Fisher Scientific, New Zealand). Traps baited with different treatment blends of HIPV compounds in five replicates were assigned in five rows, each containing treatments tested in a randomized block design. Traps were positioned 1.7 m above the ground in each trap tree, and were spaced 20 m apart in each row. The polyethylene sachets and the vial contain acetic acid were placed in the center of the sticky base in the. The first field experiment was conducted between from 1-15August 2104 in organic apple orchard (46°17′21.08″N; 119°37′0.15″W) to investigate binary and ternary blends of HIPVC and active compound obtained in previous study (El-Sayed et al. 2016). The loadings of the five HIPV blends were prepared as follows: 1) 100 mg benzyl nitrile + 3mL acetic acid; 2) 100 mg 2-phenyethanol + 3 mL acetic acid; 3) 100 mg phenylacetaldehyde + 3 mL acetic acid; 4) 100 mg benzyl nitrile, 100 mg 2-phenyethanol + 3mL acetic acid; 5) 100 mg benzyl nitrile, 100 mg phenylacetaldehyde + 3mL acetic acid; 6) 100 mg 2-phenyethanol, 100 mg phenylacetaldehyde + 3mL acetic acid; 7) 100 mg benzyl nitrile, 100 mg 2-phenyethanol, 100 mg phenylacetaldehyde + 3mL acetic acid. A trap baited with 3mL of acetic acid alone and a blank lure were used as controls. The second field experiment was conducted from 30 August to 2 September 2104 in a conventional apple orchard (46°42′28.52″N; 120°39′35.33″W) testing the same treatments and using the same protocol.

### Data Analysis

The variance of mean captures obtained with each compound or each blend of compounds was stabilized using √ (x + 1) of counts for tests of significance of treatments using ANOVA. Significantly different treatment means were identified using Tukey test was used to identify significantly different means (SAS Institute Inc. 1998).

## Results

### Volatiles emitted by uninfested and PLR larval infested apple trees

Analysis of the headspace of uninfested and infested apple trees indicated qualitative differences in odour profiles. We identified a total of 10 compounds in the headspace of uninfested apple trees, and 15 compounds in the headspace of the infested apple trees (Table 1). Five compounds (benzyl alcohol, benzyl nitrile, phenylacetaldehyde, indole, and (*E*)-nerolidol) were present only in the headspace of infested apple trees. In addition to the qualitative differences, infestation of apple trees with PLR larvae resulted in a change in the ratio of the compounds emitted from infested apple seedlings (Table 1). In contrast to our previous study (El-Sayed et al. 2016), 2-phenylethanol was not detected in the headspace of infested apple trees. Benzyl alcohol, indole, and (*E*)-nerolidol were not tested in this study because of the lack of activity in previous studies (El-Sayed et al. 2016, El-Sayed unpublished).

### Attraction of herbivore and predator to HIPVC

The composition of the HIPVC blends significantly affected the number of PLR males and females (Treatment, *F_7,32_* = 2.9, *P* < 0.02 for male, and *F_7,32_* = 6.7, *P* < 0.001 for female) and adult lacewings, *Chrysoperla carnea* (Stephens) captured (Treatment, *F_2,28_* = 4.4, *P* < 0.03). The numbers of male moths captured in traps baited with blends containing either or both of benzyl nitrile, 2-phenylethanol plus acetic acid were significantly higher than traps baited with acetic acid alone (Fig. 1). Similarly, the largest numbers of females were caught in traps baited with binary, ternary, and quaternary blends containing either or both of benzyl nitrile, 2-phenylethanol + acetic acid (Fig. 1). In contrast to herbivores, the largest number of adult lacewings were caught in traps baited with the ternary blend containing benzyl nitrile, 2-phenylethanol plus acetic acid (Fig. 2). The lowest catch of lacewings was in traps baited with benzyl nitrile plus acetic acid, while no insects were caught in traps baited with acetic acid alone (Fig. 2).

**Figure 1.**
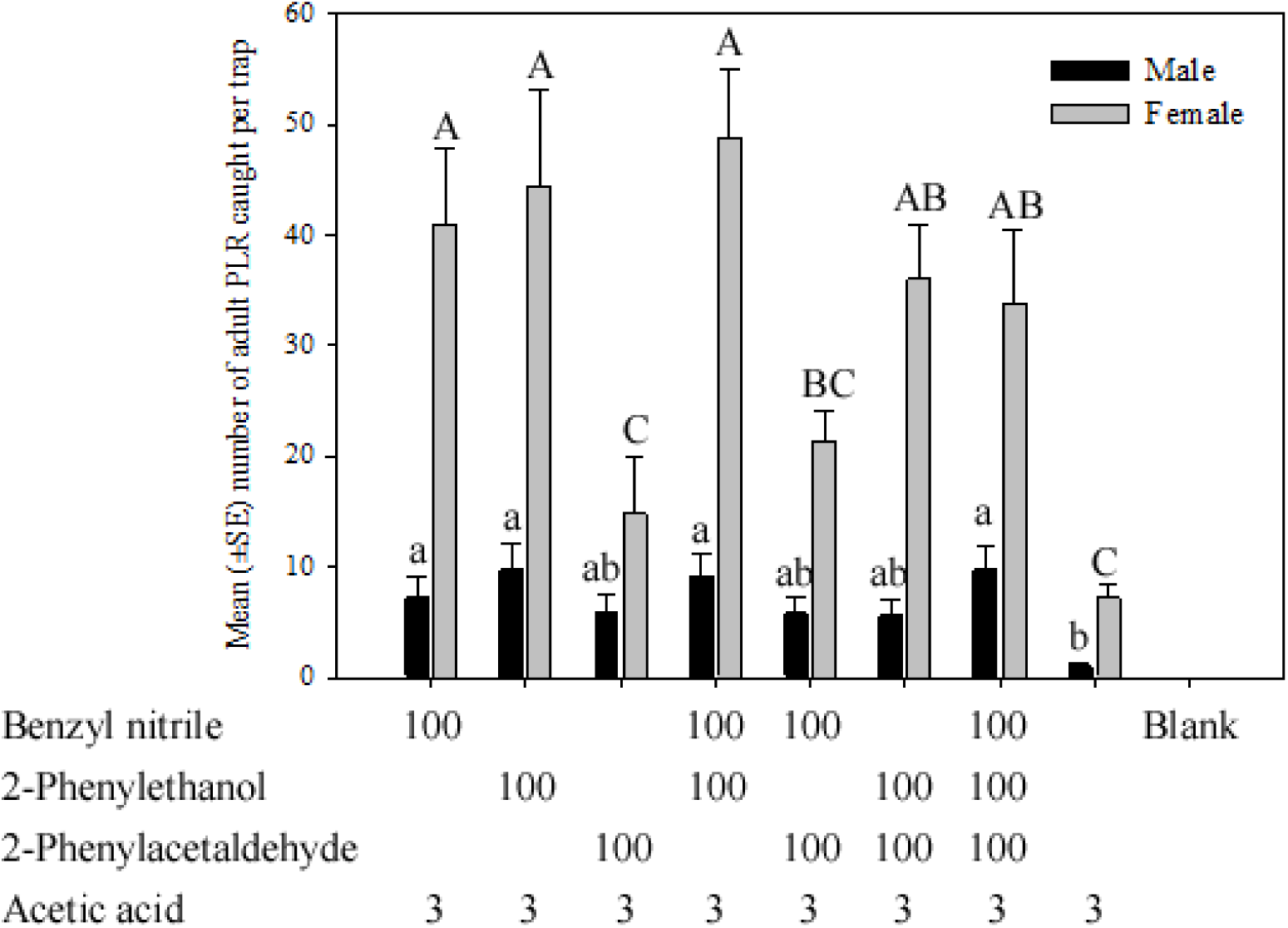
Mean (±SE) of the total number of adult male and female *Pandemis pyrusana* (Tortricidae) caught in traps baited with binary, ternary and quaternary blends of HIPVC. Loadings of the first three compounds are in mg, whereas the loading of acetic acid is in mL. Treatments labelled with the same case letters are not significantly different (*P* > 0.05, Tukey test).

**Figure 2.**
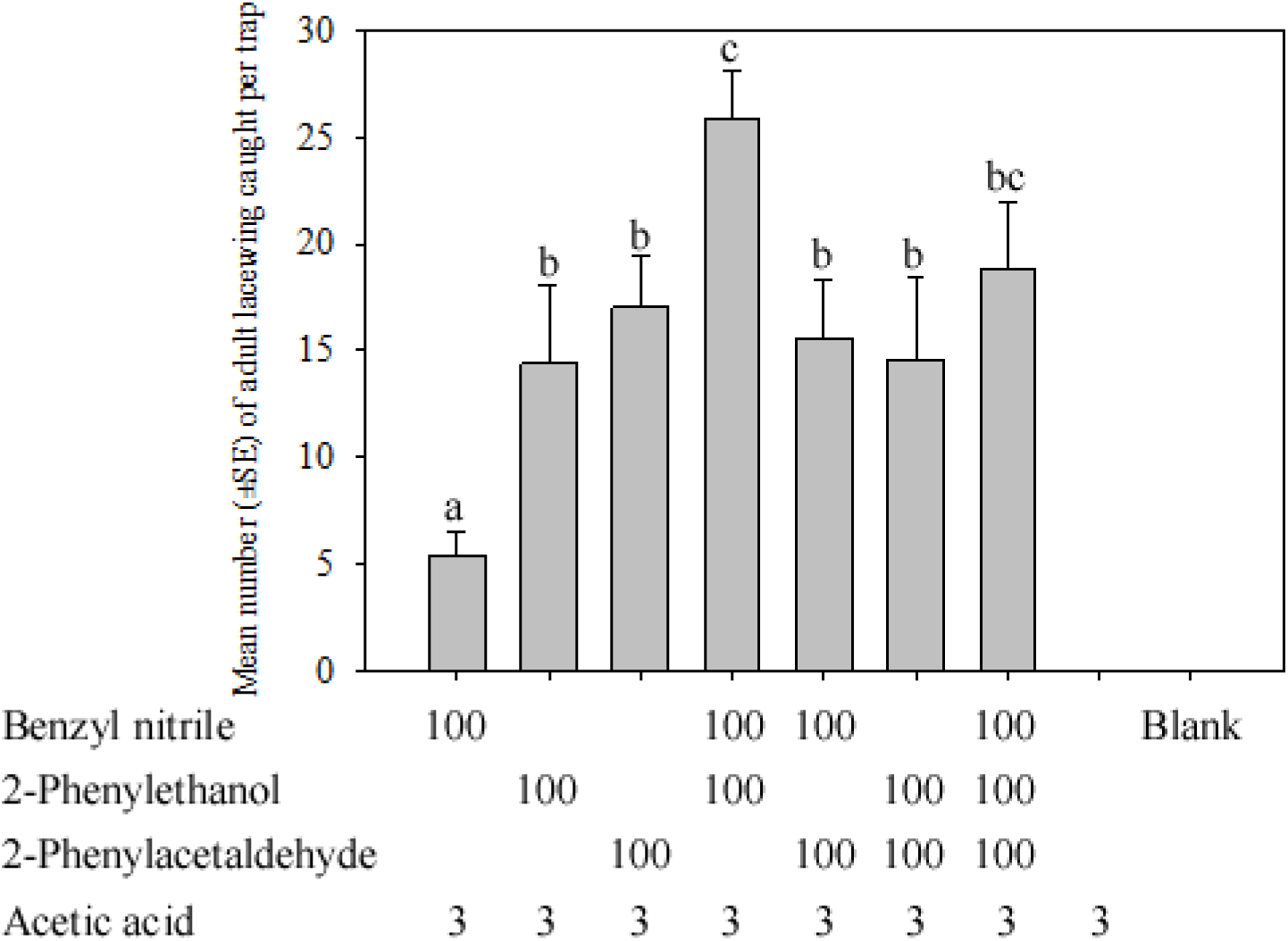
Mean (±SE) of the total number of adult *Chrysoperla carnea* (Neuroptera) caught in traps baited with binary, ternary and quaternary blends of HIPVC. Loadings of the first three compounds are in mg, whereas the loading of acetic acid is in mL. Treatments labelled with the same case letters are not significantly different (*P* > 0.05, Tukey test).

## Discussion

Qualitative and quantitative differences in the emission of volatile organic compounds (VOCs) were observed between apple trees infested with PLR larvae and uninfested apple trees. Infested plants uniquely produce the five VOCs benzyl alcohol, benzyl nitrile, phenylacetaldehyde, indole, and (*E*)-nerolidol. All five compounds were minor, where terpenes were the most dominant compounds in the headspace of infested apple trees. In contrast to apple trees infested with OBLR and ESBM larvae, 2-phenylethanol was not observed in the headspace of apple trees infested with PLR. Acetic acid and acetic anhydride were reported in the headspace of apple tree infested with LBAM larvae (El-Sayed et al. 2016). In this study we could not verify the presence of these two compounds because of the limitations of the technique used to collect headspace volatiles. The selection of the four compounds benzyl nitrile, 2-phenylethanol, phenylacetaldehyde, and acetic acid was based on chemical analysis conducted in this work and in our previous study (El-Sayed et al. 2016).

A binary blend of benzyl nitrile + acetic acid, or 2-phenylethanol + acetic acid was the most attractive blend to PLR males and females. A combination of these two compounds + acetic acid did not result in a significant increase in the number of males and females captured. Similarly, these two blends were the most attractive for other leafrollers including ESBM and OBLR (El-Sayed et al. 2016). PLR responds to 2-phenylethanol in spite of the fact it was not present in the headspace of infested apple trees. This could be due to two reasons: 1) PLR is a polyphagous species that feeds on many hosts, and 2-phenylethanol could be produced when PLR larvae feed on other host plants than apple; and 2) Ability of PLR adults to eavesdrop on other leafroller species that share the same host plants as OBLR and ESBM. The catch of females in traps baited with HIPV compounds was four to five-fold higher than the catches of males. This could reflect the sex ratio of the PLR population during the trial, in our previous work the sex ratio was almost even (El-Sayed et al. 2016).Previous work with LBAM indicated that the majority of captured females were mated, suggesting that these females were seeking ovipositor sites (El-Sayed et al. 2016)

The general predator, the common green lacewing, *C. carnea* was attracted to the same HIPVC that attracted con-specific adult herbivores including benzyl nitrile, 2-phenylethanol, and acetic acid. In contrast to conspecific herbivores, a ternary blend of benzyl nitrile + 2-phenylethanol + acetic acid attracted the largest number of *C. carnea.* However *C. carnea* responded also to a binary blend of benzyl nitrile + acetic acid, or 2-phenylethanol + acetic acid, which demonstrates the unspecific response of *C. carnea* to HIPVC. Our results show that HIPVC identified from infested apple trees attracted both a general predator and conspecific adult herbivores thus confirming the double-edged sword effect hypothesis in the system studied here. Similarly, *Nicotiana attenuate* (Torr. ex S. Watson) plants use the same defensive chemical signal to attract both herbivores and a general predator (Halitschke et al. 2008). In this study, the generalist predator, *Geocoris pallens* (Stål) showed unspecific response to HIPVC emitted from infested *N. attenuate* plants (Halitschke et al. 2008).

The attractive nature of the HIPVC to the generalist predators or specialized parasitoids has been well documented in many plant-insect systems (Dicke and Baldwin 2010). However, there is a discrepancy in the literature regarding the response of adult herbivories to plants infested with conspecific larvae. In some cases, adult herbivores were attracted to plants infested with larvae (Rojas 1999; Sun et al. 2014; Anderson and Alborn 1999; Shiojiri and Takabayashi 2003) while other herbivores were deterred by infested plants (De Moraes et al. 2001; Signoretti et al. 2012; Reisenman et al. 2013). The discrepancy reported in several plan-insect systems in relation to the response of herbivores to HIPVC could simply be due to due to experimental problems in demonstrating the beneficial effects of HIPVC as an indirect plant defence strategy. In addition, most of these studies were designed to investigate individual response in multi-tropic systems under laboratory conditions rather than focusing on the collective responses under natural conditions. The potency of the binary blends in attracting conspecific adults in our study raises an important question, what are the advantages for conspecific adult herbivores to be attracted to infested plants? Infested plants might be more favourable oviposition sites because plant resistance may be much lower and survival higher than with healthy uninfested plants (Halitschke et al. 2008; Anderson and Alborn 1999). The attraction of males to HIPVC could be due either to the presence of females in the traps or that the probability of encountering a mate would be higher at infested plants compared to uninfested plants.

The three compounds (benzyl alcohol, benzyl nitrile, and phenylacetaldehyde) identified in infested leaves are well known floral volatiles and this will result in the attraction of wide range of heterospecific herbivores as well as flower visitors (El-Sayed 2016). In addition, the attraction of conspecific adult herbivores to HIPVC would raise an important question regarding the indirect defence function of these compounds. Therefore, an integrative approach to characterize all advantages and disadvantages of these compounds for the plants is still required.

The finding of this study demonstrates two opposite functions of HIPVC under natural conditions. We anticipate this might have a negative impact on the application of these compounds in pest management of these important herbivores. To alleviate this negative effect, it might be possible to include other factors (e.g. other compounds and application during specific periods) that would allow these compounds to specifically target herbivores while preventing the attraction of natural enemies. On the other hand, it might be possible to target the predator by inclusion of other compounds that are inhibitory to herbivores.

This study is the first to identify HIPVC that directly attract both herbivore and a natural enemy under natural conditions. This result indicates that the paradigm regarding tritrophic interactions may not be as well understood as presumed previously. Significantly large number of PLR females and adult *C. carnea* were caught in traps baited with HIPVC. Such numbers have never been reported before for any tortricid female. This finding, together with our recently published results with other species (El-Sayed et al. 2016) indicates that this phenomena is widespread among leaf-feeding moths.

## Acknowledgments

This work was supported by the New Zealand Institute for Plant and Food Research Ltd. with core funds from the Ministry of Business Innovation and Employment. Ashraf El-Sayed acknowledges the receipt of a fellowship from the OECD Co-operative Research Programme: Biological Resource Management for Sustainable Agricultural Systems in 2015. The authors would like to acknowledge of the technical assistance provided by USDA co-operators during the field trials.

